# Cardiac myosin filaments are directly regulated by calcium

**DOI:** 10.1101/2022.02.19.481172

**Authors:** Weikang Ma, Henry Gong, Lin Qi, Suman Nag, Thomas C. Irving

**Affiliations:** BioCAT, Department of Biological Sciences, Illinois Institute of Technology, Chicago, IL, USA; Department of Biochemistry, Bristol Myers Squibb (TM), Brisbane, CA 94005

## Abstract

Contraction of cardiac and skeletal muscle is initiated by calcium (Ca^2+^) binding to regulatory proteins on actin-containing thin filaments. During muscle contraction, structural changes in Ca^2+^-dependent thin filament allow the binding of myosin motors to drive muscle contraction^1^. The dynamic switching between the resting *off* states and the active *on* states of myosin is also critical in regulating muscle contractility^2–4^. However, the molecular switch on the myosin-containing thick filament that drives this process is not understood. Here we show that cardiac thick filaments are directly Ca^2+^-regulated. We find that Ca^2+^ progressively moves the myosin heads from ordered *off* states close to the thick filament backbone to disordered *on* states closer to the thin filaments. This Ca^2+^-dependent structural shift of myosin is accompanied by a biochemically defined transition from the inactive super-relaxed state(s) to the active disordered relaxed state(s)^3^. Furthermore, we find that this Ca^2+^-mediated molecular switching is an intrinsic property of cardiac myosin but only when assembled into thick filaments. This novel concept of Ca^2+^ as a regulatory modulator of the thick filament provides a fresh perspective on cardiac muscle regulation, which may be particularly valuable for devising restorative treatments for pathologies altering the Ca^2+^ sensitivity of the sarcomere.

---

Calcium (Ca^2+^) signaling coordinates many different intracellular processes in plant, animal, and human physiology^5^. Muscle contraction is one of those biological processes regulated by Ca^2+^ and is propelled by the sliding of actin-containing thin filaments along myosin-containing thick filaments in the sarcomere^6^. For more than fifty years, it has been generally accepted that Ca^2+^ binds to the troponin complex on the thin filament to initiate and regulate this process^7^. Briefly, under resting conditions, the myosin-binding sites on the thin filament are blocked by tropomyosin, thus preventing muscle contraction. Upon receiving the contraction signals, Ca^2+^ entering the cytosol binds to troponin-C on the thin filament, triggering a series of conformational changes that ultimately unblock the myosin-binding site on actin by physically moving the tropomyosin^1,8^.

However, this Ca^2+^-mediated thin filament-based mechanism does not address how the thick filament is activated. Muscle myosins from invertebrates such as scallop can be activated by direct Ca^2+^ binding to the essential light chain (ELC)^9^. Tarantula skeletal muscle and vertebrate smooth muscle activate their thick filaments through Ca^2+^ binding to calmodulin, which activates myosin light chain kinase (MLCK), which in turn phosphorylates the myosin’s regulatory light chain (RLC)^10,11^. Linari et al. proposed a strain-dependent thick filament activation model (“mechanosensing”) in vertebrate skeletal muscle^2^ and later expanded this mechanism to rodent cardiac muscle^12,13^. In this mechanosensing model, early activation of a small number of *on* myosin heads generates sufficient force to strain the thick filament backbone inducing a “structural transition” that releases myosin heads from inactive *off* state(s), presumably the energy-sparing super-relaxed (SRX) state(s)^14^, to a disordered-relaxed (DRX) *on* state(s) that increases the proportion of myosin heads competent to bind actin and generate force^2,13^. It has been shown, however, that stretching alone cannot account for the full number of myosin heads that transition from the *off*-to-*on* states^15–17^, suggesting that thick filament strain cannot be the only trigger for the *off*-to-*on* structural transition resulting in the release of myosin heads from the thick filament backbone. One unexplored candidate for this additional trigger in contracting muscle would be Ca^2+^, the primary difference between active and resting muscle.

## Small-angle X-ray diffraction studies

Here we decoupled Ca^2+^-mediated thin filament-based regulation from thick filament-based regulation, using an excess concentration (100 μM) of a small-molecule inhibitor of the thin filament system. This inhibitor works by enhancing the Ca^2+^ off rate from troponin while retaining Ca^2+^ binding to troponin (Fig. S1 and Fig. S2). Examining the structural transitions using small-angle X-ray diffraction from permeabilized porcine cardiac myocardium at low Ca^2+^ concentration (pCa 8) in the presence of the inhibitor shows characteristic relaxed X-ray diffraction patterns similar to those reported previously^18,19^. With increasing Ca^2+^ concentration, the intensities of all the myosin-based reflections (M3, M6, MLLs in Fig. 1a) become weaker. The intensities of the sixth-order actin-based layer line (ALL6, Fig. 1a) are relatively stable throughout the experiment indicating that the changes in the thick filament myosin-based reflections are due to the direct effects of Ca^2+^-binding and not a result of radiation damage or by strong binding of cross-bridges to the thin filament.

**Figure 1.**
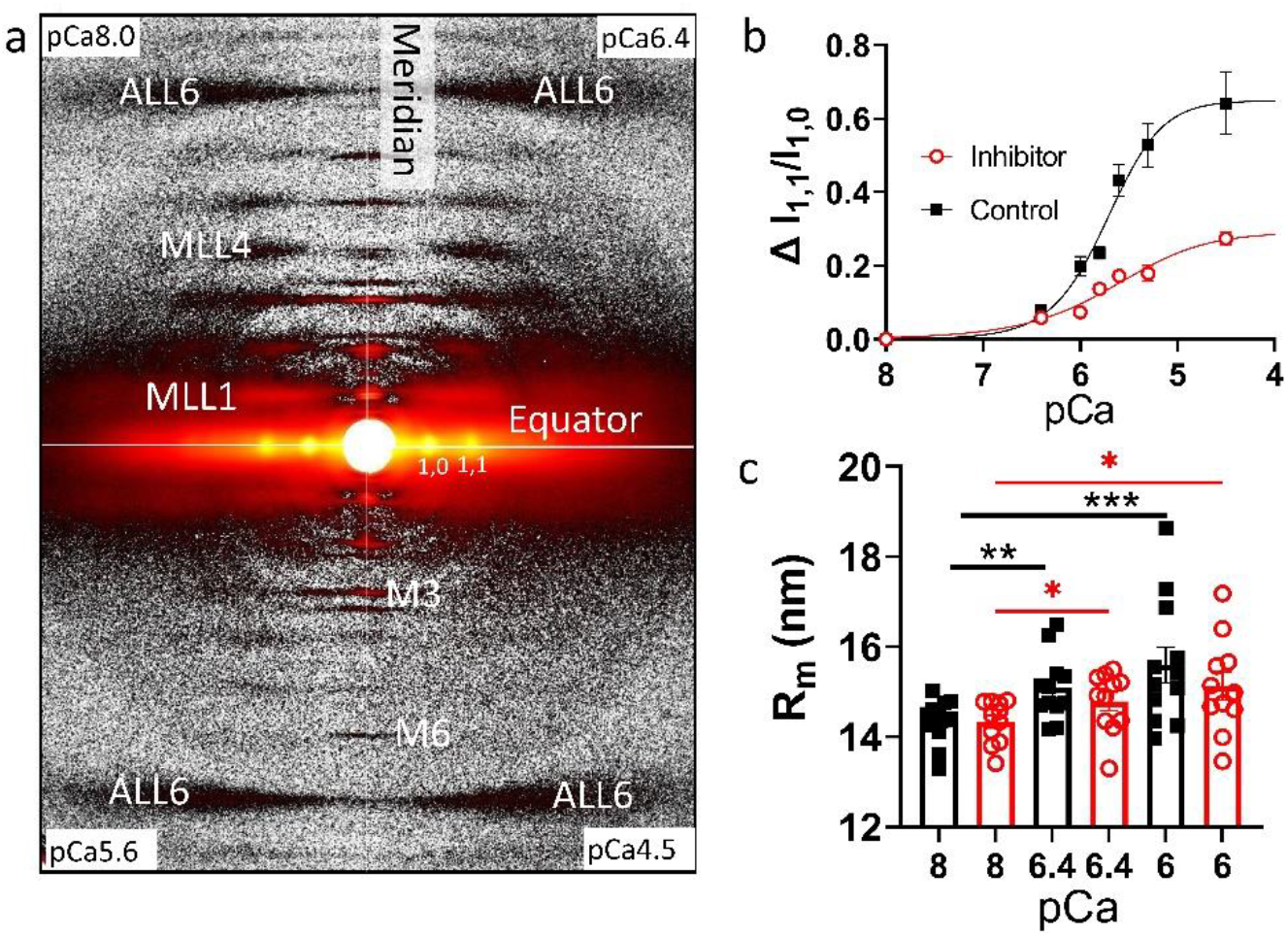
Myosin heads move radially away from the thick filament backbone. **(a)** Representative X-ray diffraction patterns from permeabilized porcine myocardium in different Ca^2+^ in the absence of force. (**b)** Change of equatorial intensity ratio (Δ I_1,1_/I_1,0_) at different Ca^2+^concentrations in the presence (red) and absence of inhibitor (black) **(c)** The radius of the average mass of myosin heads (R_m_) at different Ca^2+^ concentrations in the presence (red) and absence of inhibitor (black). Ca^2+^ induces a shift in the distribution of myosin heads away from the thick filament backbone towards the thin filaments.

## Radial movement of myosin heads

The equatorial intensity ratio (I_1,1_/I_1,0_) is indicative of the proximity of myosin heads to actin in relaxed muscle and is closely correlated to the number of force-producing cross-bridges in activated muscle^20–23^. The change of I_1,1_/I_1,0_ (Δ I_1,1_/I_1,0_) vs. pCa shows a sigmoidal shape with a pCa_50_ of 5.7 (5.5 to 5.9 for 95% CI) for the control group (n = 12). Surprisingly in the inhibitor group (n=11) where active contractions are completely eliminated, I_1,1_/I_1,0_ also progressively increases with increasing Ca^2+^ concentration (Fig. 1b), with a pCa_50_ of 5.6 (3.5 to 5.9 for 95% CI), although the amplitudes of the change are smaller compared to the control group. The radius of the center of mass of the cross-bridges (R_m_), which directly measures the proximity of helically ordered myosin heads to the thick filament backbone^21,24^, increases from 14.34 ± 0.14 nm at pCa 8 to 15.59 ± 0.4 nm at pCa 6 in the control group., R_m_ increases similarly in the presence of the inhibitor (14.34 ± 0.14 at pCa 8, to 15.13 ± 0.32 nm at pCa 6). Both the ΔI_1,1_/I_1,0_ and R_m_ data indicate that with increasing Ca^2+^ concentration, myosin heads move radially away from the thick filament backbone under conditions where they cannot bind to thin filaments, suggesting a regulatory role of Ca^2+^ directly on the thick filament.

## Structural *off*-to-*on* transitions

In a resting muscle, the majority of myosin heads are quasi-helically ordered on the surface of the thick filament, where these *off*-state myosin heads produce the myosin-based layer line reflections. Myosin heads lose their helical order when turned *on* to participate in contraction^25^. The intensity of the first-order myosin-based layer line (I_MLL1_) and the third-order myosin-based meridional reflection (I_M3_), both of which correlate with the ordering of myosin heads^26^, decreases progressively in the presence of increasing Ca^2+^. The I_MLL1_ reflection changes (Fig. 2a) with a pCa_50_ of 6.05 (5.91 to 6.19 for 95% CI) and 6.09 (5.98 to 6.19 for 95% CI) for the inhibitor and control group, respectively, whereas the I_M3_ reflection changes (Fig. 2b) with a pCa_50_ of 6.17 (6.03 to 6.33 for 95% CI) and 6.36 (6.24 to 6.51 for 95% CI) for these two experimental groups respectively. Compared to pCa 8, the I_MLL1_ and I_M3_ intensities at pCa 4.5 decrease to 27 ± 2.5 % and 35 ± 1.9 % respectively in the inhibitor group, whereas it decreases to 18 ± 1.6 % and 26 ± 2.4 % in the control group (inset in Fig. 2a and b).

**Figure 2.**
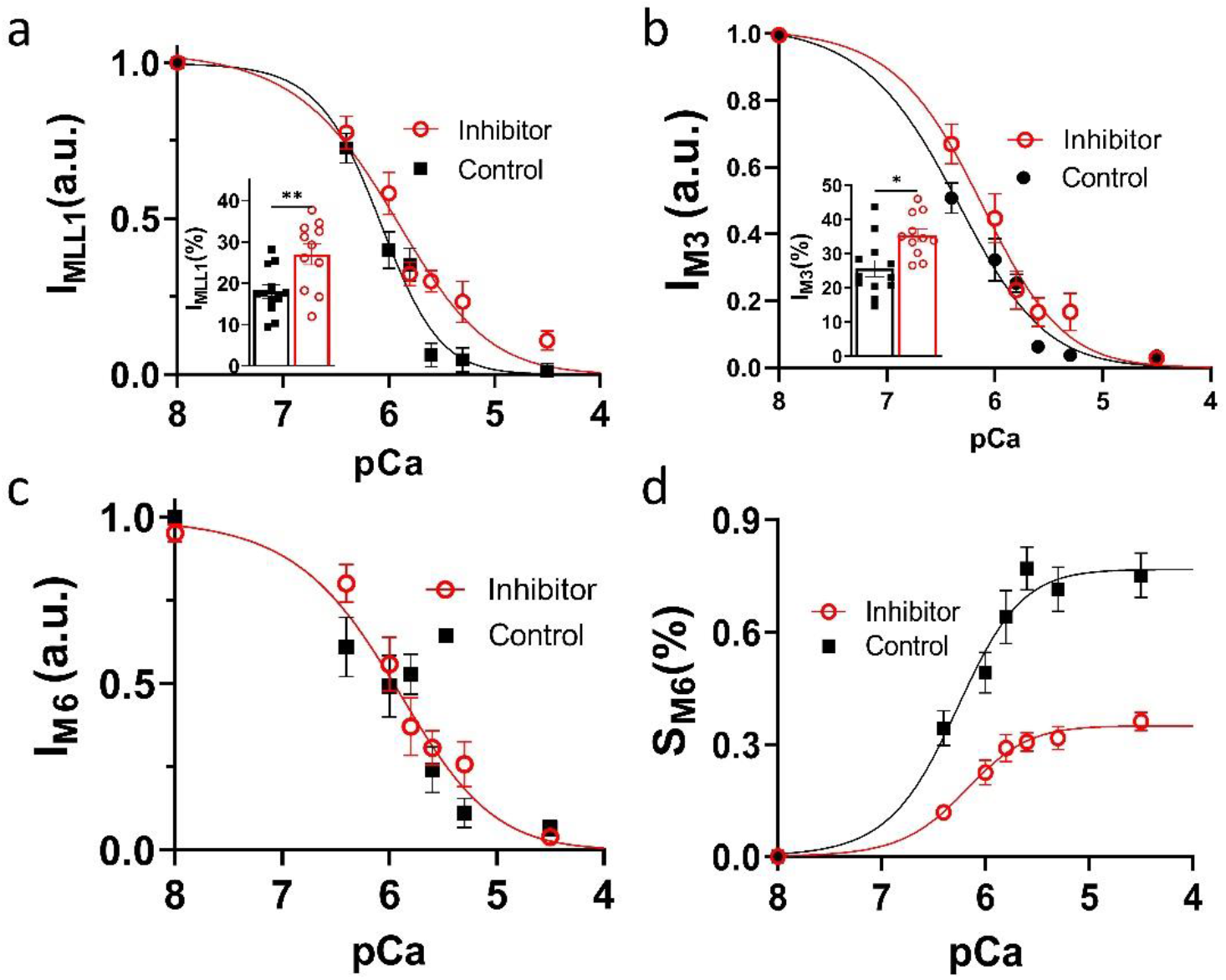
Thick filament structure changes in the presence of Ca^2+^. The intensity of first-order myosin-based layerline (**a**) and the third-order of myosin-based meridional reflection (**b**) in different concentrations of Ca^2+^ in the presence (red) and absence of inhibitor (black). The intensity (**c**) and spacing (**d**) of the sixth-order of myosin-based meridional reflection in different Ca^2+^ concentrations in the presence (red) and absence of inhibitor (black). Ca^2+^ reduces the proportion of myosin heads in ordered states on the thick filament and induces structural changes in the thick filament backbone.

Both I_MLL1_ and I_M3_ data show that myosin heads lose their helical ordering in the presence of Ca^2+^, which strongly indicates switching from the ordered *off* states to the disordered *on* states in cardiac muscle thick filaments. The diminished I_MLL1_ and I_M3_ intensities at pCa 4.5 and the leftward shift of the intensity decay curve in the control group indicate that active tension can also activate the residual myosin heads on the thick filament backbone as expected from the mechanosensing thick filament activation mechanism. However, the majority of the loss of ordering in the myosin heads can be accounted for by Ca^2+^-mediated activation of the thick filament in the absence of cross-bridge forces.

The decay of the intensity of the sixth-order myosin-based meridional reflection (I_M6_), which arises primarily from thick filament backbone^26^, with increasing Ca^2+^ concentration is indistinguishable between the inhibitor and control groups (*p* =0.6) with a similar pCa_50_ of 5.96 (5.78 to 6.13 for 95% CI). At this time, there are no obvious mechanistic explanations for a decay in I_M6_ in the presence of Ca^2+^ and the absence of force. Since I_M6_ arises primarily from the thick filament backbone, the decrease of I_M6_ in the presence of Ca^2+^ suggests a structural change in the thick filament backbone induced by Ca^2+^, independent of thick filament strain. The spacing of the M6 reflection (S_M6_), which reports the periodicity of the thick filament backbone, increases to 0.73 ± 0.06% at the fully activated state (pCa 4.5) in the control group and 0.36 ± 0.02 % in the inhibitor group where active force is absent. While the larger S_M6_ changes in the control group are caused by both the release of *off* states myosin heads and the increase in strain on the thick filament by active contraction^27^, the increase in S_M6_ in the absence of active force suggests that an, at least partial, *off*-to-*on* transition of the thick filament can occur solely in response to Ca^2+^.

## SRX to DRX transitions

Given our observation of Ca^2+^-induced *off*-to-*on* transitions of the thick filament in the absence of active force, and inspired by previous work suggesting that there might be direct effects of Ca^2+^ on thick filaments^28–30^, we explored whether these structural transitions can be translated into functional alterations. The thin filament inhibitor we used in this study has fluorescence in the 400-500 nm range, making performing conventional SRX/DRX assays on muscle tissue impractical. In order to test our hypothesis, since our results suggest that myosin or myosin filaments are the direct targets of Ca^2+^ binding, we turned our focus to reconstituted cardiac synthetic thick filaments (STF), which can faithfully capture SRX/DRX transitions under various conditions^31^.

The basal myosin SRX population in STF system at pCa 8 is 15 ± 5 %, which progressively decreases (p<0.01) to 3 ± 2 % (pCa_50_ of 5.5; 5.4 to 5.7 for 95% CI) at pCa 4 (Fig. 3a (normalized data), solid black symbol) with no considerable change in the ATPase cycling rate of the SRX states (n=5). This destabilization of the SRX states, which presumably populates myosin in some DRX states, along with an increase in the cycling rates of the DRX population, leads to an overall Ca^2+^-dependent increase (p<0.01) in the steady-state basal myosin ATPase activity from 0.03 ± 0.01 s^-1^ at pCa 8 to 0.09 ± 0.02 s^-1^ at pCa 4 with a pCa_50_ of 6.1 (5.9 to 6.2 for 95% CI) (Fig. 3a (normalized data), solid grey symbol) (n=3). Further investigation with the soluble S1-subfragment of the myosin molecule (Fig. 3a (normalized data), open symbols) and full-length myosin at high ionic strengths (150 mM KCl) (Fig. 3b) that do not fully form filamentous thick filaments demonstrates a loss of this Ca^2+^-dependent modulation, suggesting that the Ca^2+-^binding mechanism may not be intrinsic to myosin S1 or isolated full-length myosin but rather involve only myosins forming filamentous structures. This result is further supported by a lack of Ca^2+^-dependent activation of the ATPase in different soluble double-headed heavy meromyosin (HMM) constructs containing variable lengths of the S2-subfragment (Fig. S3) but lacks the light meromyosin (LMM) domain that can assemble myosin into bipolar thick filaments.

**Figure 3.**
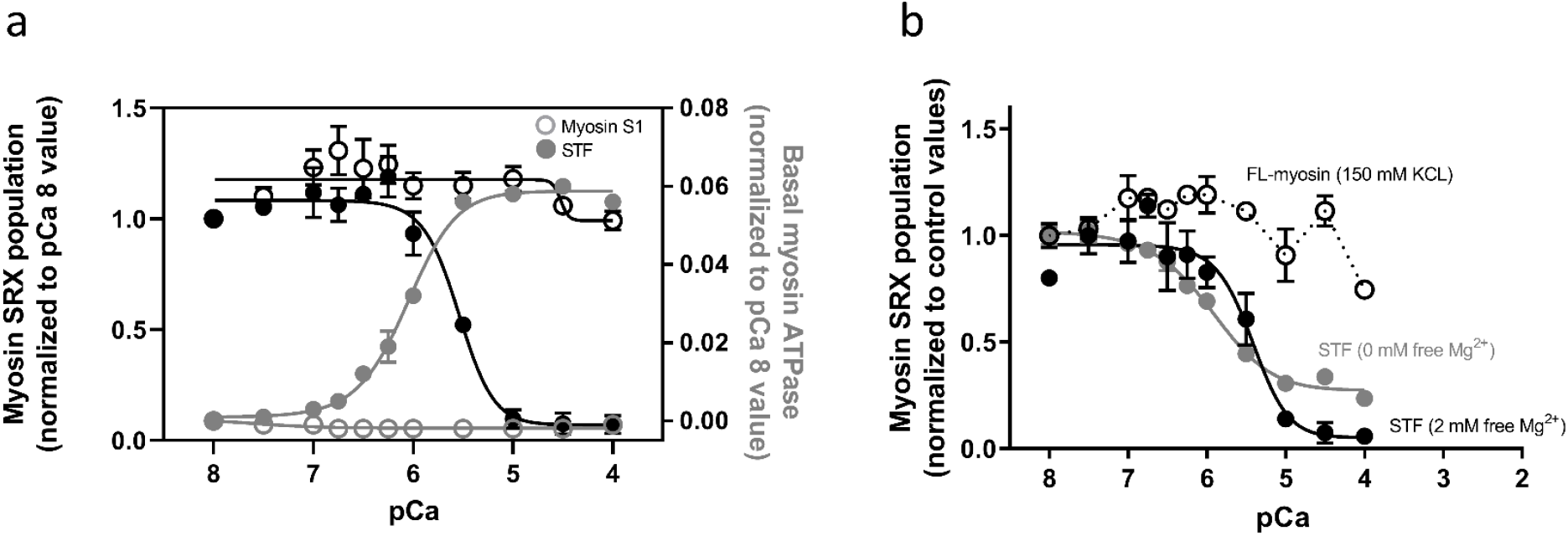
The biochemical SRX states of myosin in reconstituted synthetic thick filaments (STF) is modulated by Ca^2+^. (**a**) Normalized myosin SRX population (expressed as a fraction of the initial %SRX at pCa 8; black symbols) and normalized basal myosin ATPase activity (expressed as [ATPase value - ATPase value at pCa 8]; grey symbols) in synthetic thick filaments (STF) reconstituted from bovine cardiac full-length myosin (solid symbols) and bovine cardiac myosin S1-subfragment (open symbols) at different Ca^2+^ concentrations. (**b**) Normalized myosin SRX population in STF formed from full-length myosin at 30 mM KCL (0 mM free Mg^2+^; dark grey and 2 mM free Mg^2+^; black) and 150 mM KCl (open circles) and (solid symbols) at different Ca^2+^ concentrations. Ca^2+^ destabilizes the biochemical myosin SRX state (s) only when assembled into thick filaments.

**Figure 4.**
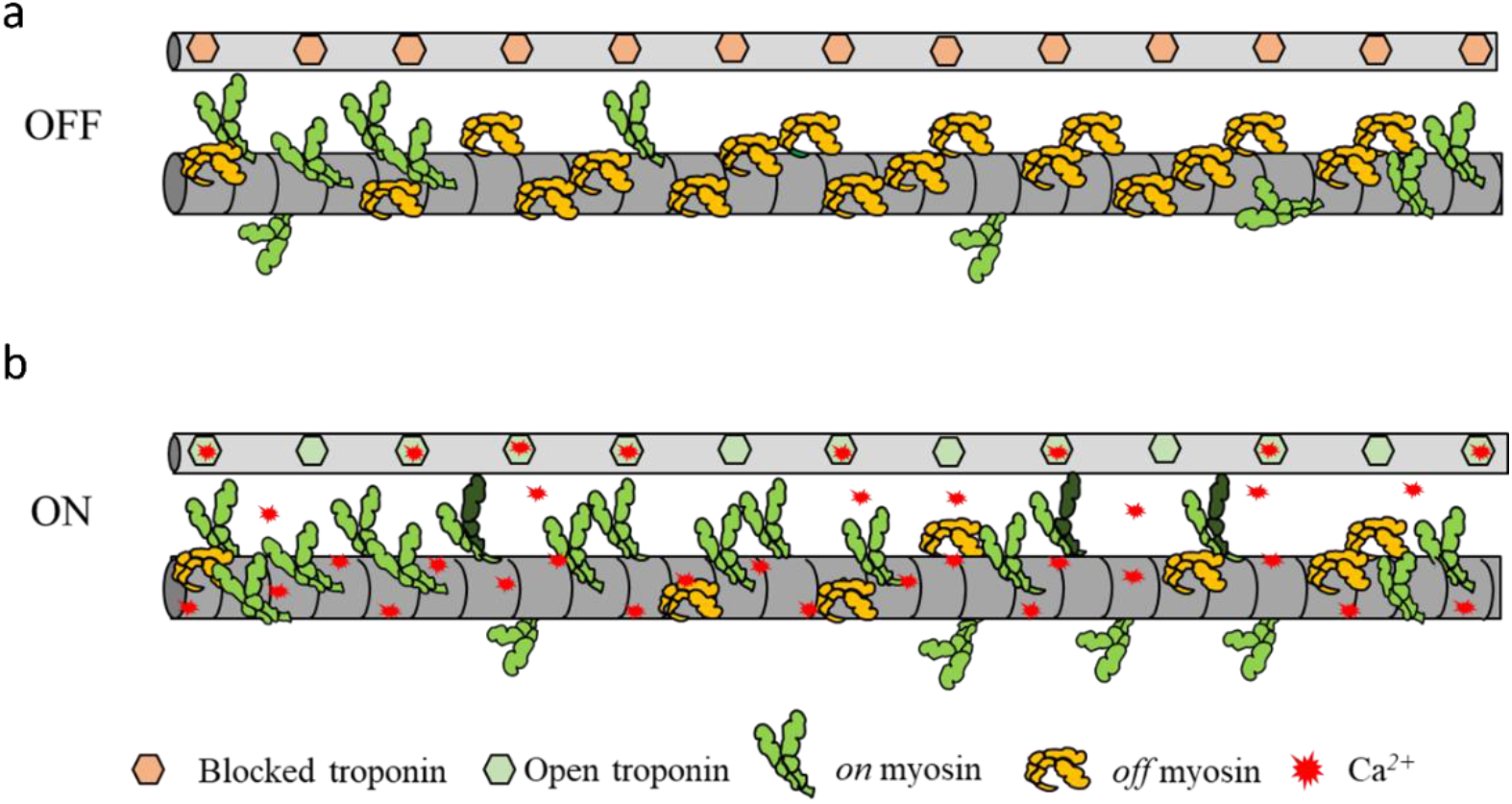
Ca^2+^-mediated dual filament regulation mechanism in cardiac muscle. A schematic cartoon of the cardiac thin (thin grey bar) and thick filament (thick grey bar) (not drawn to scale) showing the organization of the regulated thin filament and myosin. (**a**) In the absence of Ca^2+^, the resting muscle has the thin and thick filaments in the deactivated states (OFF). (**b**) Ca^2+^ (red) can independently bind to the thick and thin filament and activate them simultaneously. Ca^2+^-mediated regulation of the thick filament destabilizes the myosin heads from the *off* states (orange heads) to the *on* states (green heads), which allows the swinging of the S2-subfragment that facilitates myosin heads bind to actin. Further Ca^2+^-mediated activation of the thin filament results in active myosin cross-bridges (dark green heads) that may further destabilize myosin from the *off* states.

There is an EF-hand motif in the myosin’s RLC domain that has been shown to bind magnesium (Mg^2+^) under relaxed conditions and is increasingly occupied by Ca^2+^ as its concentration increases during muscle contraction^32^. We next study the Ca^2+^-dependent effect in the absence and presence of saturating amounts of free Mg^2+^ bound to the RLC and find that the Ca^2+^-dependent destabilization of the myosin SRX is unaltered under these two conditions, suggesting that the RLC motif is not the primary calcium transducer responsible for *off*-to-*on* transitions in the thick filament. The primary effect of Ca^2+^ on the RLC regarding thick filament *off*-to-*on* transitions appears to be indirect through MLCK phosphorylation^33^, which is not present in our experimental systems. Altogether, these data suggest that this Ca^2+^-dependent modulation of myosin is a feature that is unique to myosins assembled into thick filaments.

## Ca^2+^-mediated dual-filament regulation

In the X-ray study of permeabilized tissue, although the thin filament is shut down by the small-molecule inhibitor, other sarcomeric proteins such as myosin-binding protein C and titin are present and known to bind to Ca^2+ 34^. However, the biochemical studies with cardiac synthetic thick filament reconstituted from purified full-length myosin, showing that Ca^2+^ can bind and destabilize the SRX states of myosin, in the absence of these accessory proteins, strongly indicates that a Ca^2+^-dependent switch is an intrinsic property of myosin. At this time, the Ca^2+^-binding site(s) on the thick filament has not been identified, nor can we exclude other, less specific mechanisms. Since Ca^2+^ only turns myosin *on* when they are in filaments, and we observe structural transitions in the thick filament backbone upon Ca^2+^ binding, we strongly suspect that the Ca^2+^ binding to myosin as packed in bipolar filaments may relieve a head-backbone interaction that holds myosin heads in *off* states at diastolic Ca^2+^ concentrations.

The results presented above lead to a novel concept of a Ca^2+^-mediated dual-filament regulation model in cardiac muscle. At low Ca^2+^ concentration during diastole, both the thick and thin filaments are in *off* states. Ca^2+^ is released into the cytosol upon a nervous stimulation signal, binds to both thin and thick filaments, and allows the heart muscle to contract. The Ca^2+^ concentration can regulate the level of activation of both thin and thick filaments for them to work synergistically. The uptake of Ca^2+^ after contraction will deactivate both thick and thin filaments. For example, a recent modeling study hypothesized a Ca^2+^-dependent parked state(s) that could explain both activation and relaxation in twitches in myocardium, providing more realistic relaxation rates, resting tensions, and myosin cross-bridge detachment rate than in other current models^4^. Notably, the Ca^2+^-mediated thick filament structural transitions shown in Figures 1 and 2 and the functional transitions shown in Figure 3 have a pCa_50_ in the range of 5.5-5.9, similar to the pCa_50_ of the force pCa curve (5.91± 0.1, Fig. S1), well within the physiological range of systolic Ca^2+^ concentrations.

Additionally, there is a growing understanding that increased mitochondrial Ca^2+^ can augment ATP production^35^. These results indicate that evolution might have found an effective way to modulate cardiac muscle activation and relaxation by synchronizing both the thin and thick filament of the sarcomere as well as the energy supply by a single messenger, Ca^2+^. Imbalance in any of these components (Ca^2+^ flux, thin-or thick-filament Ca^2+^ sensitivity, and the roles of titin and MyBP-C) could disrupt the exquisite equilibrium of the system, leading to compensatory effects that cause long-term damage. We have, at this time, many unanswered questions. For example, is Ca^2+^-mediated thick filament activation fast enough to happen on a beat-to-beat basis? Is this activation sensitive to sarcomere length and relevant to length-dependent activation? These questions provide motivation for future experiments.

## Conclusion

Ca^2+^ plays a central role in cardiac muscle function, but its role has been traditionally attributed to thin filament-based regulation inside the sarcomere. Our discovery of a direct Ca^2+^-mediated destabilization of the *off* states myosin on cardiac thick filaments warrants a reconstruction of previous understandings of the roles of Ca^2+^, including Ca^2+^ sensitivity and Ca^2+^ handling, in cardiac muscle in health and disease. Over the last decade, muscle therapeutics have focused on directly targeting the sarcomeric proteins for a range of cardiac pathologies^36^, including hypertrophic cardiomyopathy, which is hypothesized to be due to increased liberation of myosin heads from the sequestered *off* states and thereby increasing force-producing cross-bridges^37^, leading to hypercontractility and diastolic impairment. This Ca^2+^-mediated thick-filament sensing mechanism for the regulation of force generation in cardiac muscle may have broader implications for existing therapeutics and the development of future sarcomere-based drug modalities. Additionally, this Ca^2+^-mediated regulation of thick filaments, observed here in cardiac muscle, may turn out to be a fundamental component of all human skeletal systems, which opens the possibility of new therapeutic approaches for many congenital myopathies caused by sarcomeric protein mutations. More generally, insofar a significant fraction of the ATPase activity of the skeletal muscle myosin is to maintain temperature homeostasis in homeothermic organisms^38^, this new role of calcium, as a driver of myosin ATP consumption, may provide a new perspective on the energetic costs of thermo-regulation.

## Acknowledgments

This research used resources of the Advanced Photon Source, a U.S. Department of Energy (DOE) Office of Science User Facility operated for the DOE Office of Science by Argonne National Laboratory under Contract No. DE-AC02-06CH11357. This project is supported by grant P30 GM138395 from the National Institute of General Medical Sciences of the National Institutes of Health. The content is solely the authors’ responsibility and does not necessarily reflect the official views of the National Institute of General Medical Sciences or the National Institutes of Health. We thank Ariana Combs and Stephen Langer of the Leinwand laboratory for generating the recombinant human cardiac 2-hep and 25-hep HMM samples. We thank Sampath Gollapudi of Bristol Myers Squibb for his contribution to the biochemical SI data on the inhibitor. We thank James A. Spudich of Stanford University, Michael Geeves of the University of Kent, Michael Regnier of the University of Washington, Srboljub Mijailovich of the Illinois Institute of Technology, and Robert S. McDowell of Bristol Myers Squibb (formerly MyoKardia) for providing helpful comments to the manuscript.

## Contributions

W.M, S.N, T.I designed the experiment; W.M, H.G, S.N performed the research; W.M, L.Q, S.N analyzed the data; W.M, S.N, T.I wrote the manuscript. All authors approved the final version of the manuscript.

## Ethics declarations

S.N is an employee and owns shares in Bristol Myers Squibb (formerly MyoKardia). T.I provides consulting and collaborative research studies to Edgewise Therapeutics, but such work is unrelated to the content of this article.

## Methods

Detailed methods along with additional data are available in the Supplemental Material

### X-ray Diffraction on permeabilized porcine myocardium

Porcine myocardium samples were prepared as described previously^18^. Briefly, frozen porcine left ventricle wall were defrosted in skinning solution (1% Triton-X100 and 3% dextran at pH 7) at room temperature for 1h before being dissected into smaller strips. The tissues were skinned at room temperature for 3 hours. The tissues were then washed by pCa 8 solution three times, 10 min each. X-ray diffraction experiments were performed at the BioCAT beamline 18ID at the Advanced Photon Source, Argonne National Laboratory^39^. Preparations were then attached using aluminum T-clips to a hook on a force transducer and a static hook. The muscles were incubated in a customized chamber, and experiments were performed at 28 °C to 30 °C at a sarcomere length of 2.3 μm. The X-ray patterns were collected at seven different Ca^2+^ concentrations with and without 100 μM inhibitor on a MarCCD 165 detector (Rayonix Inc., Evanston IL) with a 1 s exposure time. The data were analyzed as described previously^18^ using data reduction programs belonging to the open-source MuscleX software package developed at BioCAT^40^.

### Biochemical measurements in reconstituted myosin constructs

β-cardiac full-length myosin from the bovine left ventricle was isolated following established methods described elsewhere^41^. Bipolar synthetic thick filaments were prepared as described previously^31^. The same full-length myosin stock was used to prepare S1-subfragment (chymotryptic S1 without the RLC) using methods described elsewhere^41^. Myosin ATPase activity was measured at 23°C on a plate-based reader using an enzymatically coupled assay ^31^. Estimations of the SRX population of myosins were performed using a single ATP turnover kinetic protocol with a fluorescent 2’/3’-O-(N-Methylanthraniloyl) (mant)-ATP probe chased with excess unlabeled ATP^31^. The fluorescence decay profiles were fit to a bi-exponential function to estimate four parameters corresponding to fast and slow phases –the normalized % amplitude and the observed ATP turnover rate of each phase^31^. The fast and the slow phases correspond to the myosin activity in the DRX and SRX states, respectively.

### Statistics

Statistical analyses were performed using GraphPad Prism 9 (Graphpad Software). The results are given as mean ± SEM. One-way repeated measures ANOVA with the Geisser-Greenhouse correction and Tukey’s multiple comparisons test with individual variances computed for each comparison is performed on bar graphs in Fig 1 and Fig 2. The relative changes versus pCa curves were fit to a modified Hill equation to calculate the pCa_50_ values reported in the article. Table S1 reports fit parameters for all experiments. Symbols on figures: *: p<0.05,**: p<0.01, ***: p<0.001. For Fig. 3, each experiment was repeated at least twice with a minimum of two replicates per experiment, and a two-tailed student’s t-test was used to differentiate the changes in parameters among groups (p<0.01).

## Notes

### Competing Interest Statement

The authors have declared no competing interest.

