## Supplemental information for "Cardiac myosin filaments are directly regulated by calcium"

### Supplemental Material

### Methods

#### Muscle Sample Preparation

Permeabilized tissues were prepared as described previously[1]. Briefly, frozen wild type porcine (n = 2) left ventricle wall (about 1cm$^3$) was defrosted in skinning solution (91 mM $K^+$-propionate, 3.5 mM $MgCl_2$, 0.16 mM $CaCl_2$, 7 mM EGTA, 2.5 mM $Na_2$ATP, 15 mM Creatine phosphate, 20 mM Imidazole, 30 mM BDM, 1% Triton-X100 and 3% Dextran at pH 7) at room temperature before dissecting into smaller strips (~1 cm long and 2-3 mm wide). The tissues were permeabilized at room temperature for 3 hours. The tissues were then washed three times, 10 min each in pCa 8 solution (91mM $K^+$-propionate, 3.5mM $MgCl_2$, 0.16 mM $CaCl_2$, 7 mM EGTA, 2.5 mM $Na_2$ATP, 15 mM Creatine phosphate, 20 mM Imidazole and 3% Dextran at pH 7). Well-aligned tissues were further dissected into preparations of 4 mm length and a diameter of ~200 µm prior to attaching aluminum T-clips to both ends.

#### Cardiac Protein Purification

Bovine cardiac actin, tropomyosin, and troponin complex were purified following modified methods previously published[2]. Actin was stored at -80˚C as G-actin and polymerized fresh for each day of experiments by adding 50 mM KCl and 2 mM $MgCl_2$ to the actin-containing buffer. The regulated thin filament (RTF) system was reconstituted using bovine cardiac actin: bovine cardiac tropomyosin: bovine cardiac troponin complex (1:1:1 troponin-C: troponin-T: troponin-I) in a 7:1:1 ratio.

Human β-cardiac 2-hep and 25-hep heavy meromyosin (HMM) was purified using methods described elsewhere[3]. These HMM cDNA constructs consist of a truncated version of MYH7 (residues 1-855), corresponding to S1-subfragment and the first 2 heptad (14 amino acids) or 25 heptad repeats (175 amino acids) of S2-subfragment for the 2-hep and 25-hep HMM respectively, followed by a GCN4 leucine zipper to ensure dimerization. This is further linked to a flexible GSG (Gly-Ser-Gly) linker, then a GFP moiety followed by another GSG linker, and finally ending with an 8-residue (RGSIDTWV) PDZ binding peptide.

#### Calcium dissociation from the thin filament

$Ca^{2+}$ dissociation rate from the RTF system was measured in a stopped-flow instrument (KinTek Model AutoSF-120) using transient kinetic measurements. Briefly, the RTFs (with a final concentration of 7 μM actin, 2 μM tropomyosin, and 2 μM troponin) were preincubated with ~1 μM $Ca^{2+}$ (pCa 6) and was rapidly mixed with a fluorescent calcium-chelator, Quin-2, of a final concentration of 50 μM in a buffer containing 20 mM Tris-HCL (pH 7.4), 10 mM KCl, 3 mM $MgCl_2$, and 1 mM DTT. Experiments were performed at 25°C by exciting Quin-2 at 310 nm, and monitoring the emission at 450 nm.

**X-ray Diffraction**

X-ray diffraction experiments were performed at the BioCAT beamline 18ID at the Advanced Photon Source, Argonne National Laboratory [4]. The X-ray beam energy was set to 12 keV (0.1033 nm wavelength) at an incident flux of ~5 $\times 10^{12}$ photons per second. The specimen to detector distance was ~ 3 m. The preparation was then attached to a hook on a force transducer (Model 402B Aurora Scientific Inc., Aurora, ON, Canada) and a static hook. The muscle was incubated in a customized chamber whose bottom was attached to a heat exchanger, so the solution was kept between 28 °C to 30 °C. For remote solution changes, the chamber was connected to a multiway valve syringe pump (Hamilton model 500). The muscles were stretched to a sarcomere length of 2.3 µm using micromanipulators attached to the hooks while monitoring light diffraction patterns from a helium-neon laser (633 nm) on a screen. The X-ray patterns were collected sequentially at seven increasing calcium concentrations (pCa 8, pCa 6.4, pCa 6, pCa 5.8, pCa 5.6, pCa 5.3, and pCa 4.5) in the absence or presence of 100 μM inhibitor on a MarCCD 165 detector (Rayonix Inc., Evanston IL) with a 1 s exposure time. To minimize radiation damage, the muscle samples were oscillated along their horizontal axes at a velocity of 1 - 2 mm/s. The irradiated areas were moved vertically after each exposure to avoid overlapping X-ray exposures. One to two patterns were collected under each condition, and reflection spacings and intensities extracted from these patterns were averaged.

**X-ray data analysis**

The data were analyzed using data reduction programs belonging to the open-source MuscleX software package developed at BioCAT[5]. The equatorial reflections were measured by the "Equator" routine in MuscleX as described previously[6]. For subsequent analyses, the four quadrants, divided by meridian and equator, of X-ray patterns were averaged together to improve the signal-to-noise ratio, and the diffuse scatterings were subtracted from the X-ray diffraction patterns with the "Quadrant Folding" routine in MuscleX. The intensities and spacings of meridional and layer line reflections were measured by the "Projection Traces" routine in MuscleX. The spacings of targeted reflections were estimated by measuring the distance from the beam center to the peak position as the centroid of the intensity in profile, considering only the top half of the diffraction peak[7]. To compare the intensities under different conditions, the measured intensities of X-ray reflections were normalized to the intensities of the sixth-order of actin-based layer line.

**Mechanical experiments.**

Force data were collected to characterize the dose-dependence of the inhibitor on the force in skinned porcine myocardium. The muscle was activated by pCa 4.5 solutions with six increasing inhibitor concentrations (0 μM, 0.5 μM, 2 μM, 5 μM, 10 μM, and 25 μM). The forces were normalized against the force generated with 0 μM inhibitor. Separate force-pCa data were collected during the X-ray experiment in the presence and absence of 100 μM inhibitor. Force measurements were normalized against the force generated at pCa 4.5 in the control experiment.

**Statistics.**

Statistical analyses were performed using GraphPad Prism 9 (Graphpad Software). The results are given as mean ± SEM. One-way repeated measures ANOVA with the Geisser-Greenhouse correction and Tukey's multiple comparisons test with individual variances computed for each comparison is performed on bar graphs in Fig 1 and Fig 2. The relative changes versus pCa curves were fit to a four-parameter modified Hill equation (minimum response + (maximum response - minimum response)/(1+$10^{h}$. (pCa50-pCa)))[8], where $pCa_{50}$ is the calcium concentration yielding a response halfway between the minimum and maximum values reported in the article. Symbols on figures: *: $p<0.05$, **: $p<0.01$, ***: $p<0.001$. For Fig. 3, each experiment was repeated at least twice with a minimum of two replicates per experiment, and a two-tailed student's t-test was used to differentiate the changes in parameters among groups ($p<0.01$).

**Results**

Actin-activated (Actin-S1) and regulated thin filament-(RTF-S1) activated ATPase activity at pCa 6 of bovine cardiac myosin subfragment S1 was measured in response to increasing concentrations of the inhibitor (Fig. S1a). The ATPase rates were normalized against the basal ATPase rates at 0 drug concentration for these two systems, 0.02±0.002 $s^{-1}$ (n=3). The compound inhibited the RTF-S1 system ($IC_{50}$ = 9 ± 3 μM), but not the Actin-S1 in a dose-dependent manner, suggesting that the mechanism of inhibition is via shutting down the RTF system and not through actin and myosin. The enhanced calcium release rate ($AC_{50}$ =3 ± 0.5 μM; Fig. S1b) can explain the ATPase inhibition without hampering calcium binding to troponin.

The inhibitor inhibited the force production of permeabilized porcine myocardium in a dose-dependent manner with an $IC_{50}$ of 3 ± 1 (Fig. S1b) (n = 6). The force dropped to zero beyond 25 μM inhibitor concentration. Permeabilized porcine myocardium produces a classic sigmoidal force-pCa curve at a sarcomere length of 2.3 μm in the absence of inhibitor with a $pCa_{50}$ of 5.91± 0.1 (Fig. S1c; black symbol) (n = 12). However, in the presence of saturating levels of the inhibitor (100 μM), no active contraction is detected at every pCa (Fig S1c; red symbols) (n = 11). 100 μM of inhibitor concentration was chosen for all X-ray diffraction experiments to ensure complete inhibition of active force.

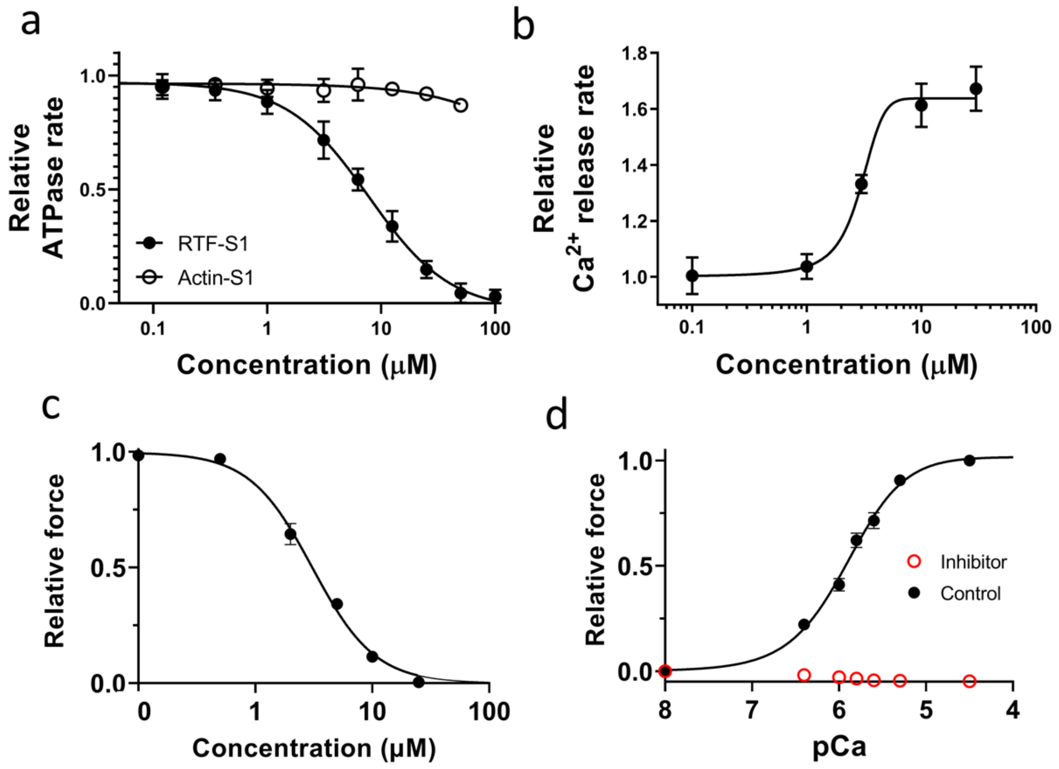


**Figure S1. Effects of the inhibitor on the chemo-mechanical activity of different sarcomere systems. (a)** The concentration-dependent steady-state ATPase activity (moles ATP used per second per mole of myosin S1 heads) of bovine cardiac myosin sub-fragment S1 with actin (open symbol) and with regulated thin filaments (RTF) at pCa 6 (closed symbols). Inhibition ($IC_{50}$ = 9 ± 3 μM ) is specific to the RTF-S1 system, implicating that the compound inhibits the ATPase activity by shutting down the RTF system and not through actin and myosin. **(b)** Concentration-dependent transient kinetics of the calcium release rate ($s^{-1}$) from the RTF system. The $AC_{50}$ of the increase in the calcium release rate is measured as 3 ± 0.5 μM. (**c**) The concentration-dependent relative maximum force of permeabilized porcine myocardium. The $IC_{50}$ of force inhibition is 3 ± 1 μM. (**d**) Relative force-pCa curve of permeabilized porcine myocardium in the absence or presence of 100 μM inhibitor. The force generated by the myocardium in the presence of the inhibitor is near-zero at all pCa values tested.

The sixth-order actin-based layer line (IALL6) intensities are relatively stable across the different calcium concentrations in control and inhibitor groups. In the control group, $I_{ALL6}$ drops to 90.55 ± 5.2 % at pCa 4.5 compared to the values at pCa8, which could be due to the disrupting of actin filament helical ordering upon cross-bridge binding. In the inhibitor group, however, $I_{ALL6}$ remains unchanged at pCa 4.5 (100.6 ± 2.4 %), as compared to the values at pCa8, indicating no structural changes in the actin filament in the presence of calcium and in the absence of active force (Fig S2). The intensity of the third-order troponin meridional reflection ($I_{Tn3}$) decreases ~ 15% in the presence of calcium in both the control group (83.5 ± 8.5 %) and the inhibitor group (87.3 ± 2.8 %), and these data indicate calcium-induced troponin structural changes are unchanged in the presence of the inhibitor, suggesting that $Ca^{2+}$ could still bind to the thin filament system in the presence of the inhibitor.

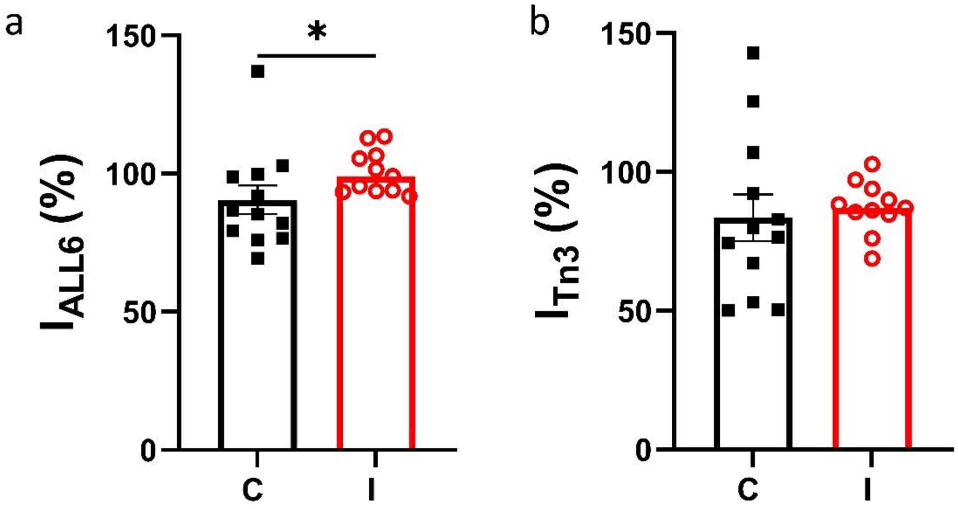


**Figure S2. Thin filament-based X-ray reflections in the presence and absence of inhibitor.** The relative intensity of the sixth-order actin-based layer line (**a**) and the third-order troponin meridional reflection (**b**) at pCa 4.5 compared to pCa 8 (C: Control; I: Inhibitor).

The steady-state basal myosin ATPase activity in STF system at pCa 8 is 0.03 ± 0.01 $s^{-1}$ which increases to 0.1 ± 0.02 $s^{-1}$ at pCa 4 with a pCa50 of 6.0 (5.9 to 6.1 for 95% CI) (Fig. S3 (normalized data)) (n=3). However, such $Ca^{2+}$-mediated ATPase activation is absent in enzymatically produced bovine cardiac HMM and S1 and recombinantly produced human cardiac 2-hep and 25-hep HMM, suggesting that the $Ca^{2+}$-mediated regulation mechanism may not be intrinsic to soluble myosin constructs but rather involve only myosins forming filamentous structures.

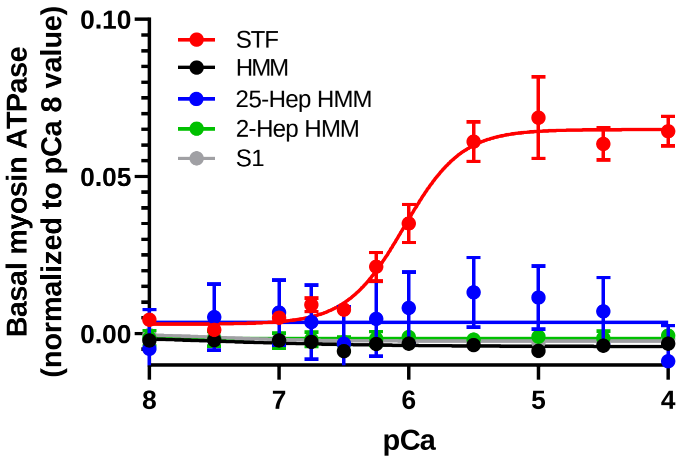

**Figure S3. The biochemical SRX states of myosin in reconstituted synthetic thick filaments (STF) is modulated by $Ca^{2+}$.** Normalized basal myosin ATPase activity (expressed as [ATPase value − ATPase value at pCa 8]; in synthetic thick filaments (STF) reconstituted from bovine cardiac full-length myosin (red), bovine cardiac HMM (black), and subfragment-S1 (grey), and in recombinant human cardiac 2-hep (green) and 25-hep HMM (blue) at different $Ca^{2+}$ concentrations.

**Table S1: Parameters obtained from fitting all datasets to modified Hill equation.**

| | Control | | Inhibitor | |
|---|---|---|---|---|
| | $pCa_{50}$ (95% CI) | Hill Slope (95% CI) | $pCa_{50}$ (95% CI) | Hill Slope (95% CI) |
| $\Delta I_{1,1}/I_{1,0}$ Fig 1b | 5.7 (5.5 to 5.9) | 0.8 (0.3 to 1.4) | 5.6 (3.5 to5.9) | 1.5 (0.9 to 2.9) |
| $I_{MLL1}$ Fig. 2a | 6.1 (6.0 to 6.2) | 1.5 (1.1 to 2.0) | 6.1 (5.9 to 6.2) | 1.4 (0. 9 to 2.2) |
| $I_{M3}$ Fig. 2b | 6.4 (6.2 to 6.5) | 1.1 (0.8 to 1.4) | 6.2 (6.0 to 6.3) | 1.3 (0.8 to 2.0) |
| $I_{M6}$ Fig. 2c | 6.0 (5.8 to 6.1) | 1.0 (0.6 to 1.4) | 6.0 (5.8 to 6.1) | 1.0 (0.6 to 1.4) |
| $S_{M6}$ Fig. 2d | 6.3 (6.1 to 6.6) | 1.4 (0.7 to 2.3) | 6.2 (6.0 to 6.4) | 1.4 (0.7 to 2.6) |
| STF SRX (2 mM $Mg^{2+}$) | 5.5 (5.4 to 5.7) Fig. 3a<br>5.4 (5.2 to 5.7) Fig. 3b | 2.4 (1.4 to 2.5) Fig 3a<br>1.8 (0.9 to 1.9) Fig 3b | | |
| STF SRX Fig. 3b (0 mM $Mg^{2+}$) | 5.9 (5.8 to 6.1) | 1.1 (0.8 to 1.5) | | |
| STF ATPase | 6.1 (6.0 to 6.1) Fig. 3a<br>6.0 (5.9 to 6.1) Fig. S3 | 1.7 (1.3 to 2.3) Fig. 3a<br>1.9 (1.4 to 3.1) Fig. S3 | | |
| Force Fig. S1d | 5.9 (5.8 to 6.0) | 1.3 (1.1 to 1.6) | | |

| | Inhibitor | |
|---|---|---|
| | $IC_{50}$ (95% CI) | Hill Slope (95% CI) |
| ATPase Actin-S1 Fig S1a | >100 μM | - |
| ATPase RTF-S1 Fig S1a | 7.8 (6.6 to 9.4) | 1.2 (1.0 to 1.5) |
| $Ca^{2+}$ release rate Fig S1b | 3.0 (2.5 to 3.5) | 0.6 (0.5 10 1.0) |
| Force Fig. S1c | 3.1 (2.7 to 3.5) | 1.6 (1.4 to 1.9) |
